# Partial selfing eliminates inbreeding depression while maintaining genetic diversity

**DOI:** 10.1101/210005

**Authors:** Ivo M. Chelo, Bruno Afonso, Sara Carvalho, Ioannis Theologidis, Christine Goy, Ania Pino-Querido, Stephen R. Proulx, Henrique Teotónio

**Affiliations:** Instituto Gulbenkian de Ciência, Apartado 14, P-2781-901 Oeiras, Portugal.; cE3c – Center for Ecology, Evolution and Environmental Changes, Faculdade de Ciências, Universidade de Lisboa, Lisboa, Portugal.; Institut de Biologie de l’École Normale Supérieure (IBENS), Inserm U1024, CNRS UMR 8197, F-75005 Paris, France.; Benaki Phytopathological Institute, Stefanou Delta 8, 14561 Kifissia, Greece.; Leibniz Research Institute for Environmental Medicine, 40225 Düsseldorf, Germany.; Department of Ecology, Evolution, and Marine Biology, University of California Santa Barbara, CA 93106, U.S.A.

**Keywords:** selfing, outcrossing, experimental evolution, deleterious recessive alleles, identity disequilibrium, genetic incompatibility, overdominance, *C. elegans*

## Abstract

Classical theory on the origin and evolution of selfing and outcrossing relies on the role of inbreeding depression created by unlinked partially-deleterious recessive alleles to predict that individuals from natural populations predominantly self or outcross. Comparative data indicates, however, that maintenance of partial selfing and outcrossing at intermediate frequencies is common in nature. In part to explain the presence of mixed reproductive modes within populations, several hypotheses regarding the evolution of inbreeding depression have been put forward based on the complex interaction of linkage and identity disequilibrium among fitness loci, together with Hill-Robertson effects. We here ask what is the genetic basis of inbreeding depression so that populations with intermediate selfing rates can eliminate it while maintain potentially adaptive genetic diversity. For this, we use experimental evolution in the nematode *C. elegans* under partial selfing and compare it to the experimental evolution of populations evolved under exclusive selfing and predominant outcrossing. We find that the ancestral risk of extinction upon enforced inbreeding by selfing is maintained when populations evolve under predominant outcrossing, but reduced when populations evolve under partial or exclusive selfing. Analysis of genome-wide single-nucleotide polymorphism (SNP) during experimental evolution and after enforced inbreeding suggests that, under partial selfing, populations were purged of unlinked deleterious recessive alleles that segregate in the ancestral population, which in turn allowed the expression of unlinked overdominant fitness loci. Taken together, these observations indicate that populations evolving under partial selfing gain the short-term benefits of selfing, in purging deleterious recessive alleles, but also the long-term benefits of outcrossing, in maintaining genetic diversity that may important for future adaptation.

## Introduction

Explaining the origin and maintenance of selfing and outcrossing within natural populations is an important goal of evolutionary biology (Darwin 1876, Fisher 1930, Stebbins 1957, Lloyd 1988, Barrett 2008). Classic theoretical models posit a key role for unlinked and deleterious partially-to fully-recessive alleles in generating inbreeding depression and in promoting the evolution of reproduction mode modifiers (Lande and Schemske 1985, Charlesworth and Charlesworth 1987, Charlesworth et al. 1990). Under selfing, more homozygotes are produced than under outcrossing and, as a consequence, fitness differences between individuals larger and selection more effective at purging deleterious recessives. In general, if mutation rates and/or their dominance effects are not strong enough to lead to extinction (Lande et al. 1994, Schultz and Lynch 1997), then selfing should evolved, otherwise outcrossing should evolve.

Comparative studies in plants and in animals have found that a number of natural populations show mixed reproduction modes, with selfing and outcrossing being frequent (Jarne and Charlesworth 1993, Goodwillie et al. 2005, Jarne and Auld 2006, Wright et al. 2013). Sexual selection or an equilibrium between selection for sex allocation and for the acquisition of resources for survival and reproduction irrespective of sex may result in partial selfing and outcrossing, e.g. (Lloyd 1976, Porcher and Lande 2005, Anthes et al. 2010, Escobar et al. 2011, Carvalho et al. 2014b, Poullet et al. 2016). Similarly, fluctuating selection on selfing and outcrossing, for example due to co-evolving pathogens or due to metapopulation dynamics, could explain the maintenance of partial selfing, e.g., (Pannell and Barrett 1998, Morran et al. 2011, Cheptou 2012, Masri et al. 2013). While these “ecological” factors are important, the acknowledgement that fitness loci are always in linkage or identity disequilibrium, due to variable recombination rates or selfing and assortative mating (Weir and Cockerham 1973), respectively – together with the interactions between genetic drift and selection (Hill-Robertson effects) – brought back into focus the special role of the genetics of inbreeding depression in the evolution of reproductive modes, e.g., (Lande et al. 1994, Winn et al. 2011, Kamran-Disfani and Agrawal 2014, Roze 2015, 2016). That unlinked overdominant loci may cause inbreeding depression has also been appreciated, e.g. (Holsinger 1988, Ziehe and Roberds 1989, Charlesworth and Charlesworth 1990, Bierne et al. 2000, Chelo and Teotónio 2013). Of particular interest here, elucidating the genetic basis of inbreeding depression may explain how partially-selfing populations retain genetic diversity of a potential adaptive nature, and, if so, whether it is independent or not of how selfing reduces effective population sizes (Pollak 1987), effective recombination (Nordborg 1997), and/or of how it influences selection on recombination modifiers (Roze and Lenormand 2005).

Experimental data directly supporting a crucial role of inbreeding depression in the origin and evolution of partial selfing is weak, given the difficulty of controlling reproduction mode and standing genetic variance for fitness in most organisms, but see, e.g., (Porcher et al. 2004, Weeks 2004, Harder and Barrett 2006, Noel et al. 2017). Evolution experiments done with the androdioecious nematode *Caenorhabditis elegans* are a singular exception (Anderson et al. 2010, Teotónio et al. 2017). *C. elegans* is a bacteriofagous androdioecious species, where hermaphrodites are capable of autonomous selfing or of outcrossing with males (Maupas 1900). Sex determination is chromosomal with hermaphrodites XX and males XØ (Nigon 1949) and, since hermaphrodites cannot mate with each other, outcrossing rates are expected to be twice the male frequency in the absence of segregation distortions (Stewart and Phillips 2002, Cutter et al. 2003, Teotónio et al. 2012). Under several demographic scenarios, hermaphrodites are limited in self-reproduction as they first produce sperm and then irreversibly switch to oogenesis (Barker 1992, Cutter 2004). Genetic manipulation of reproduction mode in *C. elegans* is possible with sex-determination mutants, and populations with standing genetic variation and domesticated to standard laboratory conditions have been obtained (Teotónio et al. 2017).

Wild isolates of *C. elegans* do not show inbreeding depression but outbreeding depression (Dolgin et al. 2007, Chelo et al. 2013a), because of a history of predominant selfing, metapopulation dynamics and linked selection (Cutter 2006, Rockman et al. 2010, Andersen et al. 2012). In the laboratory, partially-selfing isogenic wild isolates exposed to relatively mild deleterious mutagenesis maintain males at higher frequencies than isolates under natural mutation rates but they eventually go to extinction (Manoel et al. 2007). Unless wild isolates are subjected to strong deleterious mutagenesis, males and outcrossing are not favored (Morran et al. 2009); but see (Carvalho et al. 2014a) for increased male frequencies under natural mutation rates when starting from hybrid inbred lines between wild isolates. For weak to mildly deleterious mutagenesis, even at small population sizes, predominant selfing is sufficient to maintain high mean population fitness (Estes et al. 2004), and only with very strong mutagenesis will selfing populations go extinct, due to fixation of mutations, and partially-selfing to obligate outcrossing populations withstand the build-up of inbreeding depression (Morran et al. 2009, Morran et al. 2010). Here we use “strength” of mutagenesis loosely since few studies have estimated the homozygous and heterozygous selection coefficients of fitness loci and there is usually the assumption that these fitness loci are unlinked, but see (Peters et al. 2003, Manoel et al. 2007).

For lab populations starting experimental evolution with standing genetic variation, partial selfing is maintained at intermediate frequencies, in part through a balance between male reproductive success and hermaphrodite self-reproduction success (Anderson et al. 2010, Teotónio et al. 2012, Masri et al. 2013, Carvalho et al. 2014b). The role of inbreeding depression in the evolution of partial selfing in these experiments is unclear, with one study indicating that initial purging of unlinked deleterious recessive alleles is later overcome by fitness overdominance (Chelo et al. 2013a), and others suggesting that outcrossing modifiers will succeed in the long term because of increased effective recombination rates in bringing together beneficial alleles (Parrish et al. 2016); see also (Morran et al. 2009, Morran et al. 2011). Our previous work on the topic, in particular, had the design problem that selective and genetic effects were confounded when measuring inbreeding depression (Chelo et al. 2013a), by assuming that biparental and uniparental inbreeding were equivalent, cf. (Porcher and Lande 2016).

Using experimental evolution in *C. elegans* we here investigate the genetic basis of inbreeding depression and how populations where selfing rates can evolve deal with it. We first characterize the survival rates of hermaphroditic lineages upon enforced inbreeding by selfing, and compare the fertility of experimental populations with that of inbred lines derived from them. We then follow how genome-wide single-nucleotide polymorphisms (SNPs), and their associations across the genome, behave during experimental evolution and after enforced inbreeding in order to characterize the genetic basis of inbreeding depression and its evolution.

## Materials and Methods

### Ancestral population

All populations employed here are ultimately derived from a hybrid population obtained by funnel-crossing 16 wild isolates at moderate population sizes to break natural linkage disequilibrium and maintain high standing levels of genetic variation (Teotónio et al. 2012, Noble et al. In Press). This hybrid population was cultured under 4-day discrete and non-overlapping life-cycles at census population sizes at the time of reproduction of 10^4^ – and effective population sizes of about 10^3^ – to derive a 140 generation lab-adapted population (Chelo et al. 2013b, Chelo and Teotónio 2013); which is the ancestor population for all the experimental populations reported here (named A6140). During lab adaptation, worms were cultured in Petri dishes filled with NGM-lite agar (US Biological), containing NaCl at 25 mM, whose surface was covered by a lawn of HT115 *E. coli* that served as the *ad libitum* food. After lab adaptation, A6140 showed a male frequency of 45% (Theologidis et al. 2014) and abundant genetic variation [average observed heterozygosity of 0.3 at genome-wide SNPs (Chelo and Teotónio 2013) and average linkage disequilibrium decay to background levels at about 0.5cM on an F2 generation genetic scale (Noble et al. In Press)].

### Experimental evolution

The experimental evolution design has been detailed in (Theologidis et al. 2014). We mass introgressed an X-chromosome sex determination mutant [*xol-1(tm3055)*] that kills male embryos into A6140 to derive a monoecious population composed exclusively of hermaphrodites (named M00; nomenclature of the experimental populations can be found in the archived data). In parallel, we mass introgressed an autosomal sex determination recessive mutant [*fog-2(q71)*] that eliminates self-spermatogenesis into A6140 to derive a trioecious population composed of males and females, and hermaphrodites at an expected frequency of 10^−5^ (named T00). We sampled the same number of families from A6140, as those during the derivation of M00 and T00, to maintain the same amount of standing variation (this later population being named A00). Replicate populations of M00, T00 and A00 were exposed in NGM-lite media to 8 mM / generation increases in the concentration of NaCl from the first larval stage until adulthood during 35 generations, after which they were kept at 305 mM NaCl for 15 extra generations. Here we report the results of 2 replicates derived from T00 that underwent 50 generations of experimental evolution [GT150, GT250; nomenclature according to (Theologidis et al. 2014)], 2 replicates from M00 (GM150, GM350), and 3 replicates from A00 (GA150, GA250, GA450). In (Theologidis et al. 2014), we found that due to developmental time problems males are lost in high salt concentrations. In Figure 1, panel A, we re-plot from (Theologidis et al. 2014) the observed male frequencies during experimental evolution.

**Figure 1.**
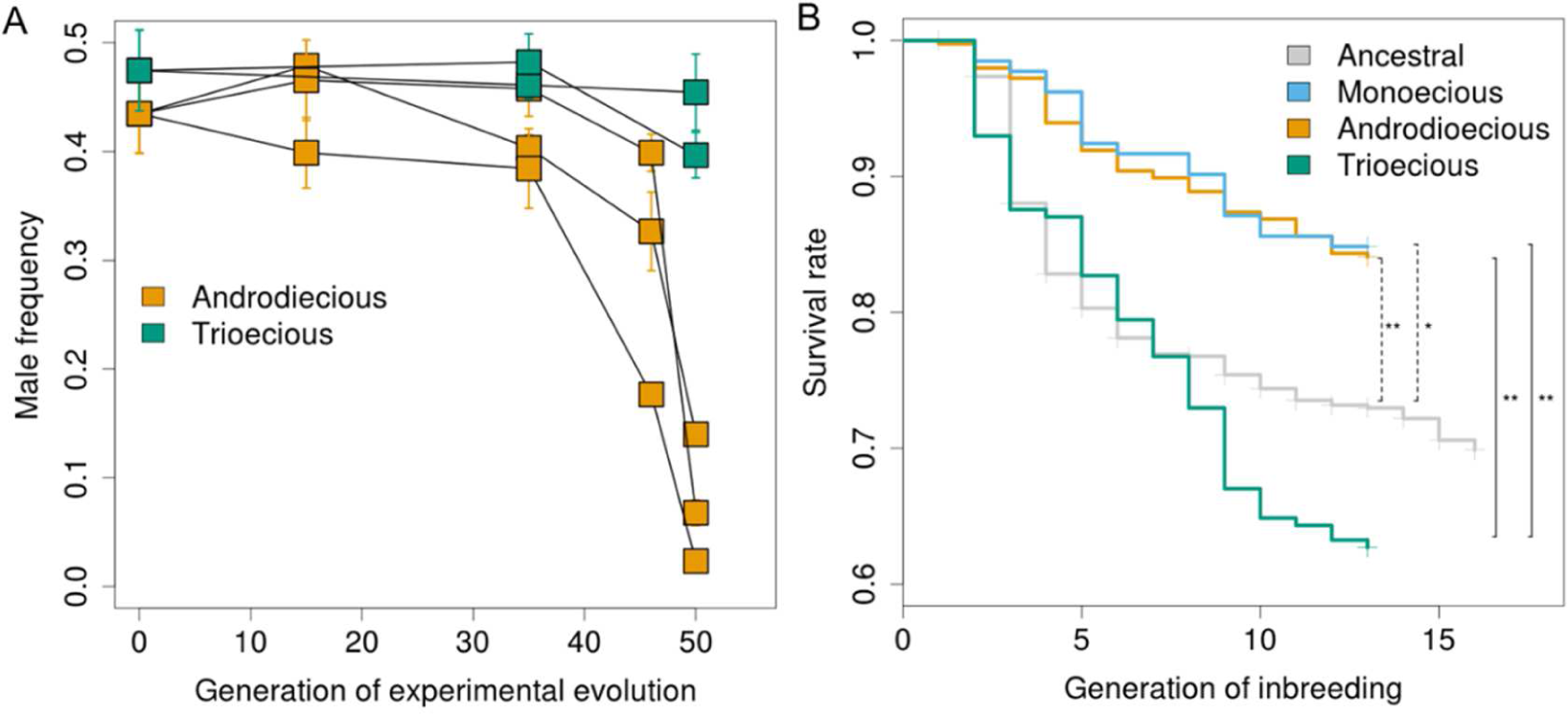
A, from (Theologidis et al. 2014), the observed male frequencies during experimental evolution under different reproduction systems. Error bars are SD obtained from estimates of different experimental blocks, each one based on three technical replicates. Monecious populations have no males by design and are not shown. B, Kaplan-Meyer estimates of survival rates (1 minus risk of lineage extinction) are shown for the laboratory adapted ancestral population (gray line) and the experimentally evolved populations. Observed survival rates per replicate population are shown in Supplementary Figure 2. Significance of the differences in risk of extinction between experimentally evolved populations and the ancestral population is shown in dashed lines: ** p-value<0.005, * p-value<0.05. Trioecious populations may show a marginal increase in extinction risk from the ancestral population (p-value=0.1). As ancestral and evolved populations were assayed at different times and blocks (see Materials and Methods), a more appropriate model only compares risk of lineage extinction among the experimentally evolved populations. Significance of the differences in risk of extinction with inbreeding between them is shown in full lines: ** p-value<0.005 (uncorrected for multiple testing). Assuming that mutations have not contributed to inbreeding depression, the patterns showed by the trioecious populations must have been due to standing variation already present in the ancestral population.

### Inbreeding depression: lineage extinction assays

Following standard protocols, we measured inbreeding depression as the survivorship of lineages upon enforced inbreeding by selfing (Charlesworth and Willis 2009). A6140, and all generation 50 populations, were revived from frozen stocks, each with >10³ individuals, and cultured for two generations. In the third generation, hermaphrodites (or possibly females from trioecious populations, see below) were handpicked at the larval immature L3-L4 stage and placed in individual wells of 12-well cell culture plates, previously filled with NGM-lite media and bacteria. During inbred line derivation NaCl was used at 25 mM, the salt condition of lab adaptation. Every 4 to 7 days, one L3-L4 offspring hermaphrodite was handpicked and placed into a fresh well, consecutively for 16 generations for A6140 or for 13 generations in all other populations. Plates were kept at 4°C to prevent growth until successful offspring production could be confirmed in the following generation. A lineage was considered extinct if reproduction from the previous generation failed twice. For A6140, inbred line derivation was done in two blocks, corresponding to the initial thawing date of the population sample. For generation 50 populations, another six separate blocks were done, with each including all populations. In each block, two experimenters derived the lines with no particular bias for the population being handled.

The risk of lineage extinction (or lineage survival rates) per inbreeding generation was assessed with Kaplan-Meyer estimators, assuming right censored data. Cox proportional hazards regression models were then done to compare the extinction risks of experimentally evolved populations against the ancestor A6140 population, with the different reproductive systems being used as predictors. Replicate populations, within each reproductive system, were considered as random factors in a mixed model. Since these estimates are confounded by potential environmental effects due to ancestral and derived populations being assayed in different blocks, we also modeled just the differences between derived populations. For all analysis, the R (R Development Core Team 2015) package *coxme* was used for computation (Therneau 2015).

By generation 50, trioecious populations still segregated females [see Figure S8B in (Theologidis et al. 2014)], and many of the hermaphrodites were heterozygous for the sex determination recessive *fog-2(q71)* mutant. There was therefore a possibility that lineage extinction was due to picking females and heterozygous hermaphrodites during the first few generations of inbreeding. We conducted numerical simulations to determine the extent of this possibility. Specifically, we started by random sampling individuals from the two trioecious experimental populations, whose genotypes at *fog-2* followed frequencies estimated by PCR in (Theologidis et al. 2014), indicating that females were at around 30% (unpublished results). For each of the resulting individuals, which were either heterozygotes or homozygotes for the selfing wild-type allele, we generated offspring (1 per each) by sampling twice from the progenitor alleles. For *fog-2(q71)* homozygote hermaphrodites, which are sterile in the absence of males, we resampled again and only once, from their respective progenitors to mimic the experimental procedure. If this process again resulted in a *fog-2(q71)* offspring being generated, then we considered that lineage to become extinct. Results from these simulations show that the observed lineage extinction in trioecious populations could not have been due to sampling effects (Supplementary Figure 1).

### Inbreeding depression: fertility assays

Assay design was previously described in (Chelo et al. 2013b, Guzella et al. 2017, Noble et al. In Press). Individual fertility was measured in all experimental populations [A6140, A00, M00, T00, and all generation 50 (G50) derived populations] and the inbred lines at 25 mM and 305 mM NaCl. Each sample was revived from frozen stocks and maintained for two generations in a standard environment to avoid confounding environmental effects (Teotónio et al. 2017). On the third generation, synchronized first larval staged L1s were seeded into regular Petri dish plates with 25 mM or 305 mM of NaCl. 48h later, L3-L4 staged hermaphrodites or females were hand-picked individually into separate wells of a 96-well cell culture plate, containing NGM with 1 uL of an O/N culture of bacteria and 25 mM or 305 mM NaCl. After single individual transfer, plates were covered with Parafilm to prevent cross-contamination and kept at 20ºC and RH 80%. 24h later, the worms were exposed to bleach solution (1 M KOH: 5% NaOCl): M9 (50:50 volumetric ratio) for 5 min, following (Teotónio et al. 2012). 200 µL of M9 liquid solution were then added to each well and rinsed 3 times. After another rinsing step, the M9 suspension with embryos, and larval and adult debris (200 µL) was transferred to another 96-well plate, which had been previously filled with 120 uL of M9. The plates were left overnight in the incubator. 24h later, plates were centrifuged for 1 min at 1800 rpm and 3 pictures were taken at 4 pixel/µm resolution with a Nikon Eclipse TE2000-S inverted microscope to provide estimates of the number of live L1s in each well. ImageJ was used for visualization. Quality control was done on the 96 well-plates and during image analysis, by identifying and removing data where bacteria were absent, worms were absent or dead at the time of bleach, males had been inadvertently hand-picked, more than 1 adult was present, and adults were still alive after the bleach hatch-off protocol.

Observation of fertility distributions revealed a large dispersion around the mean and the presence of many zero counts. Poisson and negative binomial distributions and also zero-inflated versions of the same distributions, were initially fitted independently to each population sample, to inform us on the best model to be used. Likelihood ratio tests were used for selection between nested distributions in a hierarchical fashion. R functions *glm* (stats package), *glm.nb* (MASS package) and *zeroinfl* (*pscl* package) were used for model fitting. Zero-inflated negative binomial error distributions were chosen for subsequent analyses since they provided the significantly best fit for the majority of cases (71%), followed by the negative binomial distributions (27% of cases). This was expected given that fertility, as measured in our conditions, depends on reaching reproductive maturity (Theologidis et al. 2014). We thus consider the excess of zero counts to come from those individuals not reaching sexual maturity at the time of bleach, which we refer to as the proportion of "immature individuals", and this was analyzed separately from *bona fide* fertility estimates.

Zero-inflated negative binomial distributions were then fitted to each population sample to estimate the proportion of counts in the inflated zero class. Generalized mixed effect models (function *glmmadmb* from R package *glmmADMB*), assuming negative binomial distributions for fertility data and binomial distributions for the proportion of immature individuals, were used to test for the effects of evolution and reproductive system, with experimental blocks and replicate populations as random factors.

Inbreeding depression (or enhancement) was tested as any negative (or positive) difference between experimental populations and the inbred lines derived from them, using *lsmeans* and *pairs* functions in R. Note that since L3-L4 females from the trioecious populations were picked they were unfertilized, particularly at high salt (Theologidis et al. 2014), and this assay does not fully account for fertility during experimental evolution.

### DNA collection and genotyping

Genomic DNA from A6140, M00 and each of the G50 populations was extracted from individuals with the ZyGem prepGEM Insect kit. Genomic DNA from all of the inbred lines that survived until the end of the inbreeding assays was extracted from pools of individuals using the Qiagen Blood and Tissue kit. A total of 830 SNPs distributed evenly across genetic distance, obtained by linear interpolating the F2 genetic distances of (Rockman and Kruglyak 2009), were genotyped using Sequenom methods (Bradic et al. 2011). The choice of the SNPs was based known variation between the founder wild isolates used to generate the A6140 (Noble et al. In Press). For the populations, each individual was genotyped for chromosomes I and II, III and IV, and V and X, respectively.

Quality control was done following (Chelo and Teotónio 2013), first by removing SNPs with more than 60% of missing data, then by removing individuals with more than 60% missing data, and finally by removing SNPs observed in less than 10 individuals in any given population were also removed. In the end, the resulting genotype matrix is composed of 743 SNPs. The genotyping data with different quality control criteria from A6140 and the inbred lines has been presented in (Noble et al. In Press), while that of M00 and G50 monoecious populations in (Guzella et al. 2017). Sample sizes per population, inbred line and per SNP after quality control can be found in Supplementary Table 1.

### Population genetics during experimental evolution

SNP allele diversity (*H*_*e*_) was estimated as *H*_*e*_= 2*pq* = *1-(p^2^+q^2^)*, and a fixation index as *F_is_= 1 − (H_o_/H_e_)*, with *H*_*o*_ being the observed proportion of heterozygotes (Crow and Kimura 1970). To avoid sampling biases, SNPs with allele diversity lower than 0.05 were removed prior to calculation. Linkage disequilibrium between pairs of markers was estimated as *r*^2^ = *Δ*^2^/*p*_*i*_(1-*p*_*i*_)*p*_*j*_(1-*p*_*j*_), where ∆ is the difference in genotype frequencies from that expected given the allele frequencies *p*_*i,j*_ in SNPs *i* and *j* (Weir 1996). Comparison between different populations was based on the expected *r*^2^ with genetic distance, for markers segregating between 0.05 and 0.95 of frequency: *E[r^2^]= 1/(1+4Nc)* (Sved 1971), where *c* is the genetic distance between marker pairs (Rockman and Kruglyak 2009). *N* was estimated from the data by fitting a non-linear model with function *nls* in R. Identity disequilibrium [*g2*, (David et al. 2007)] was calculated using *inbreedR* package (Stoffel et al. 2016). Identity disequilibrium quantifies at the genome-wide level the extent to which genotypes at different SNPs are correlated (Weir and Cockerham 1973). Finally, we also calculated the effective number of haplotypes (*h*_*e*_) as: 1/Σ*p*_*i*_^2^, with *p*_*i*_ being the proportion of haplotype *i* in a given replicate population or group of lines. *h*_*e*_ can be regarded as the number of haplotypes segregating in the population if all were at the same frequency (Crow and Kimura 1970). For each population, haplotypes were defined in non-overlapping windows of 10 SNPs, separately for each chromosome. We present the mean over all SNPs or haplotypes for *H*_*e*_, *F*_*is*_, *r*^2^, *g*^*2*^ and *h*_*e*_ for each population.

Differentiation of *H*_*i*_ and *F*_*is*_ between the different reproductive systems and the ancestral A6140 population was tested with *summary.lm* function in R.

### Maintenance of diversity after inbreeding

*H*_*e*_ and *h*_*e*_ metrics were calculated for the inbred lines by considering each as a diploid individual. Note that due to the sampling design, about 32 haplotypes were available from the experimental populations, whereas the final number of lines was usually above 50. For this reason, comparison of diversity estimates was done between observed data of the experimental populations and means of resampling subsets (Jackknife resampling) for the inbred lineages. After (Chelo et al. 2013a, Chelo and Teotónio 2013), and in order to provide neutral confidence intervals for genetic diversity after inbreeding, numerical simulations were done with selfing for 13 (for generation 50 derived populations) or 16 generations (for A6140). 1,000 simulations were done per population with resulting haplotypes being sampled in the same numbers as the inbred lines that were obtained from them. All chromosomes were analyzed separately. Each run started by randomly sampling with replacement phased diploids [phasing was done as described in (Chelo et al. 2013a) from the experimental population replicates in equal numbers as those of the starting inbred lineages]. Recombination was simulated by exchanging consecutive sets of alleles between the two parental haplotypes (defined as vectors of SNP alleles). We assumed complete crossover interference and map sizes of 50cM. Crossover occurred randomly between any two consecutive SNPs according to the probability given by the genetic distances between them, by joining two independent gametes to obtain the individual progeny.

## Results

### Evolution of inbreeding depression under different reproduction systems

We measured inbreeding depression first as the relative survival rate of lineages undergoing inbreeding by selfing and second as the individual fertility between ancestral and experimentally evolved populations at generation 50. Androdioecious populations quickly lost males after generation 35, having 10% of males by generation 50 (Figure 1, panel A). Trioecious populations maintained close to effective dioecy during most of experimental evolution, having above 40% of males by generation 50.

In the ancestor population only about 70% of the lineages survived for 16 generations of inbreeding (Figure 1B and Supplementary Figure 2). The monoecious and androdioecious populations evolved a lower risk of extinction, with approximately 85% of lineages surviving for 13 generations. Trioecious populations, on the other hand, evolved an increased risk of extinction compared with monoecious and androdioecious populations.

During experimental evolution fertility increased, particularly for trioecious and monoecious populations in high salt conditions (Supplementary Figure 3). However, there is little evidence that fertility of inbred hermaphrodites evolved (Supplementary Figure 4). Further, we failed to detect any fertility trade-offs between high and low salt among the surviving inbred lines (Supplementary Figure 5).

### Population genetics of SNP during experimental evolution

We followed the population genetics under different reproduction systems, by measuring the average of several genome-wide genetic diversity metrics.

The ancestor population and the trioecious populations after experimental evolution show low SNP fixation indices (*F*_*is*_; Figure 2A). In contrast with *F*_*is*_ results, SNP allele differentiation from the ancestor was similar under trioecy and androdioecy, being much lower than that found under monoecy (Figure 2B). But in line with SNP allele differentiation, trioecious and androdioecious populations maintained ancestral SNP allele diversity (*H*_*e*_), while in monoecious populations it was much reduced after experimental evolution (Figure 2C).

**Figure 2.**
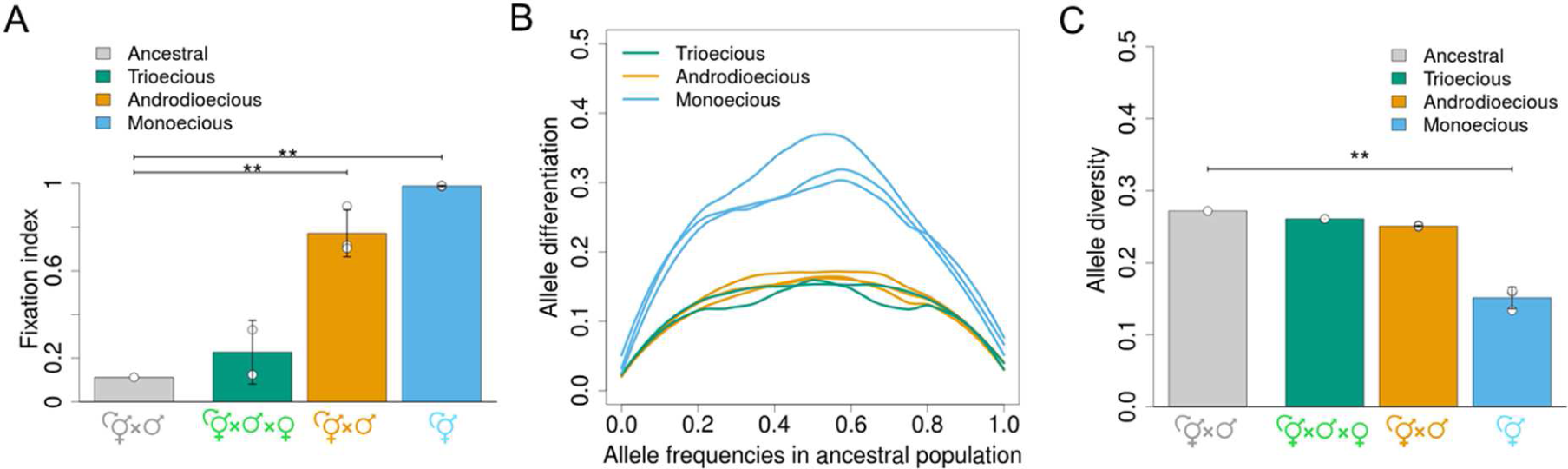
A, higher SNP fixation indexes (*F*_*is*_) are seen in monoecious and androdioecious populations than in the ancestral or trioecious populations. B, higher SNP allele frequency differentiation from the ancestral population distinguishes evolution under monoecy from that under androdioecy and trioecy. Lines are the local polynomial regression of the magnitude of change in allele frequencies on ancestral frequencies (*loess* function in R was used), shown for each replicate population. C, SNP allele diversity (*H*_*e*_) is reduced in monoecious populations. A, B, and C, asterisks show significant differences between experimentally evolved populations and the ancestral population for p-value<0.005. Allele diversity metrics are the genome-wide average of 743 SNPs calculated per replicate population. In A and C panels, error bars are standard deviation based on experimental population replicates and individual population values are shown as white dots.

We also find that trioecious and androdioecious populations show the same extent of linkage disequilibrium between pairs of SNPs (*r*^*2*^; Figure 3A). But androdioecious populations have a much higher identity disequilibrium than trioecious populations, which maintain ancestral levels (*g2*; Figure 3B). Linkage disequilibrium is very high, and identity disequilibrium complete, under monoecy.

**Figure 3.**
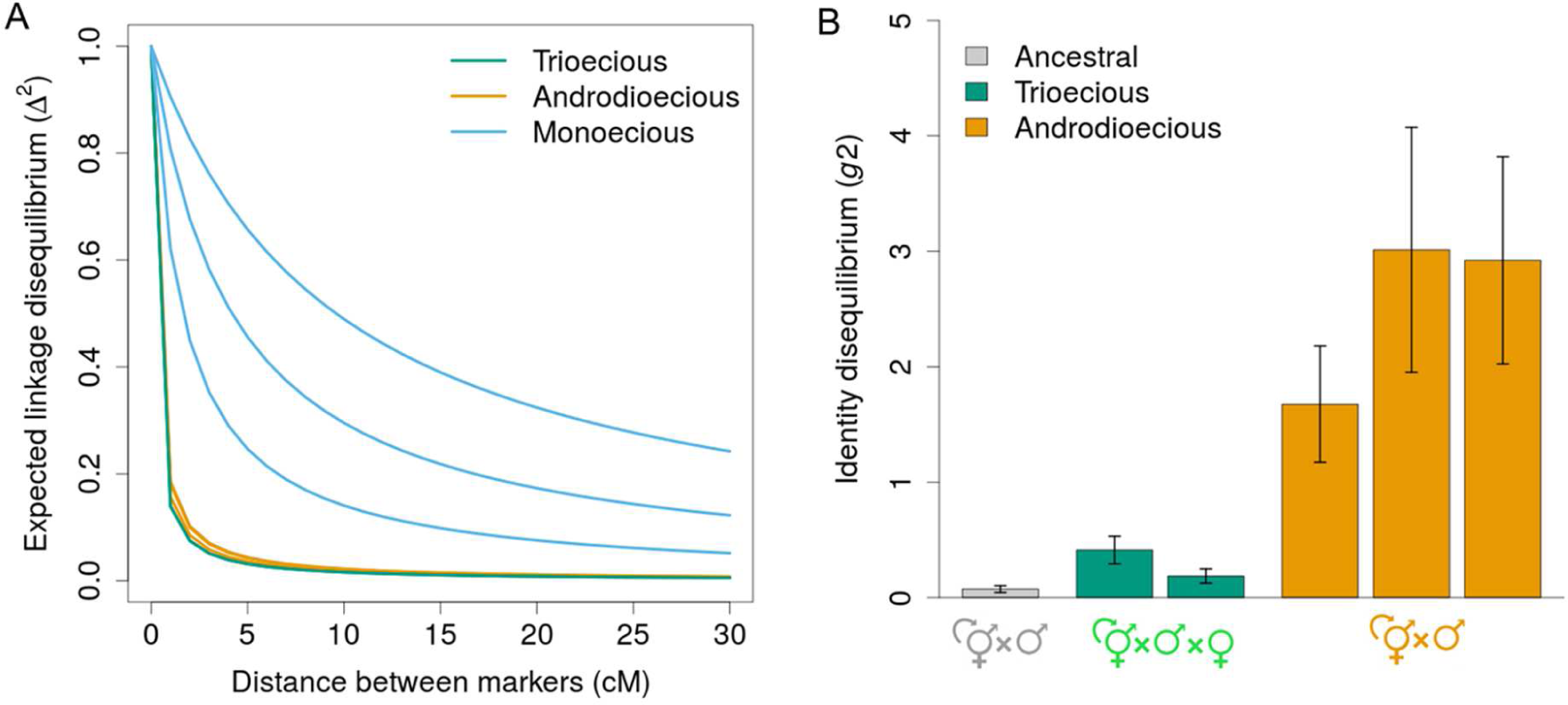
A, Decay of linkage disequilibrium (*r*^2^) with genetic distance (cM), estimated by fitting E[*r*^2^]= 1/1+*xc* with respect to *x*, is shown for each experimental population. In this formulation, *x* reflects historical effective population size (Sved 1971) and depends on the extent of selfing (Nordborg 2000). Note, however, that E[*r*^2^] refers to an expected value after an “equilibrium” state between population size, partial selfing and recombination rates has been achieved. Our experimental situation is clearly out of this equilibrium. The line referring to the ancestral population overlaps with the one from the trioecious populations and thus, it is not shown. Only linkage disequilibrium from marker pairs within chromosomes were used. B, expected identity disequilibrium (*g2*) is shown for experimental populations with exception of monoecious populations, which have no heterozygous SNPs. Reproductive system means and standard deviations are shown, together with individual population values (white dots).

### Genetic diversity after enforced inbreeding by selfing

Genome-wide SNP and haplotype diversity was also measured among the lines that survived inbreeding by selfing. For trioecious and androdioecious populations, there is a negative correlation between the directions SNP allele differentiation observed during experimental evolution with that observed after inbred line derivation (Figure 4A and B). For monoecious populations we find no such relationship (Figure 4C).

**Figure 4.**
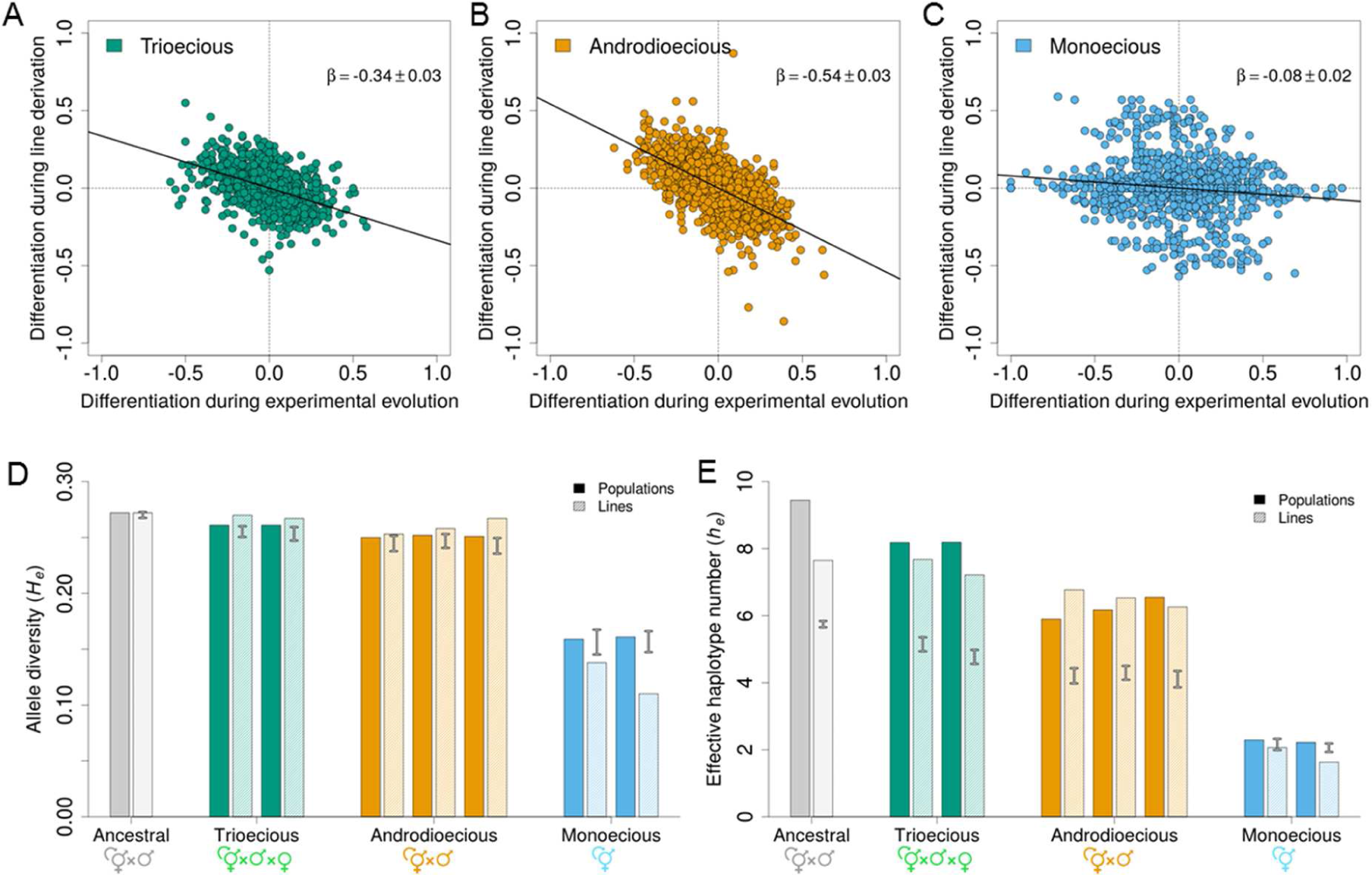
A, B and C, comparison of SNP allele differentiation from ancestral frequencies during experimental evolution and after enforced inbreeding by selfing. Data falling into the 2nd and 4th quadrants indicate cases where allele frequencies changed in opposite directions during both experimental stages. Insets show regression slopes based on all data with error being given from the 743 analyzed SNPs (all slopes are highly significant, P-value<0.001). Multilocus diversity, shown as the average SNP allele diversity (*H*_*e*_, panel D) or mean effective haplotype number (*h*_*e*_, panel E) are plotted for experimental populations and among the inbred lines derived by inbreeding. Error bars show the 95% credible intervals of the expected diversity with no selection during enforced inbreeding, as obtained with 1,000 numerical simulations (see Materials and Methods). Solid and dashed bars show, respectively, the observed diversity in experimental populations and among the inbred lines derived from each replicate population.

Trioecious and androdioecious populations show the maintenance of more SNP and haplotype diversity among the lines that survived inbreeding than that expected by sampling and chance (Figure 4D and E). Maintenance of more genetic diversity with inbreeding than expected was the ancestral condition. In contrast, monoecious populations show a larger reduction of SNP and haplotype diversity after inbreeding than expected.

## Discussion

Lineage extinction evolved to lower levels under mating systems with higher levels of selfing (Figure 1 and Supplementary Figure 2), while individual fertility did not evolve (Supplementary Figure 4), suggesting that inbreeding depression was reduced in populations that underwent selfing and that inbreeding depression was maintained in populations that outcrossed The risk of lineage extinction shown by monoecious populations provides the benchmark of the expected loss due to environmental effects (including accidental lineage death) against which androdioecious results are compared, since monoecious individuals are expected to be homozygous throughout the genome (Figure 2A). Likewise, the trioecious populations show ancestral inbreeding depression to which androdioecious can be compared, as we assume no role for new mutations during experimental evolution.

There is little evidence for heterosis. Previous work investigating several fitness components in the progeny of crosses between inbred and isogenic wild isolates, indicate that outbreeding depression, not heterosis, is the norm in *C. elegans* (Dolgin et al. 2007, Chelo et al. 2013a), as in other androdioecious *Caenorhabditis* species (Gimond et al. 2013). Consistent with the presence of outbreeding depression in our experiments, we find here that trioecious populations may have had a higher risk of lineage extinction than the ancestor population (one replicate population appears to show increased lineage extinction when compared to the ancestral, Supplementary Figure 2). Since outbreeding depression in *C. elegans* may result from the break-up of gene complexes, e.g. (Seidel et al. 2008, Noble et al. In Press), themselves evolved in nature by genetic drift or selection under predominant selfing (Cutter 2006, Rockman et al. 2010, Andersen et al. 2012), one can venture that with trioecy tightly linked natural gene complexes were further broken down because of increased effective recombination rates during experimental evolution, but see Figure S4 in (Noble et al. In Press). As we previously argued (Chelo et al. 2013a), hybridization may have brought together deleterious recessive alleles (that may have also independently accumulated in natural populations under predominant selfing), which in turn could have created inbreeding depression in our predominantly outcrossing populations (Epinat and Lenormand 2009).

Classical theory on the evolution of selfing rates suggests that if inbreeding depression is generated by unlinked deleterious partially-to fully-recessive alleles affecting a quantitative trait under stabilizing selection, then one of two stable outcomes is possible (Lande and Schemske 1985). One possibility is that high levels of selfing evolves, with populations mostly free of deleterious recessive alleles, since homozygous individuals are efficiently selected against. The other possibility is that high levels of outcrossing evolves, with populations bearing a load of deleterious recessive alleles, as selection is less efficient in purging them. Whether the population reaches a stable state of selfing or outcrossing depends on the evolutionary history of accumulation of unlinked deleterious recessives in the population being considered. Contingent on this history, modifiers of selfing rates can invade populations with the predominant alternative reproductive mode and, as a consequence, explain transitions between reproductive systems, e.g., (Charlesworth et al. 1990).

Some of our results are in line with this theory, although we are not concerned with long-term mutation accumulation but short-term segregating genetic variation in fitness loci and reproductive mode modifiers (Glemin and Ronfort 2013). As expected by the experimental evolution design and the observed evolution of selfing rates (Figure 1A), both the ancestral and trioecious populations effectively performed as fully outcrossing populations since the observed levels of heterozygosity were similar to those expected under panmixia (Figure 2). Little population genetic structure may have thus facilitated the maintenance of inbreeding depression due to deleterious recessive alleles. Conversely, because more homozygous individuals were produced by selfing in the predominantly selfing androdioecious and monoecious populations, deleterious recessives were exposed to selection, purged from the populations and inbreeding depression was eliminated during experimental evolution.

Some of our results do not fit with the classical theory, as presented above. Upon enforced inbreeding more genetic diversity was maintained under trioecy and androdioecy than expected by chance (Figure 4D and E). That selection during inbreeding could have been responsible for this pattern is demonstrated by the opposite direction most SNP differentiation occurred in comparison with experimental evolution, likely due to strong competition among progeny within individual lineages (Figure 4A and B); although the nature of this selection is not clear since we failed to find a negative correlation between, for example, fertility in the environment of experimental evolution and that employed during enforced inbreeding.

Two genetic mechanisms can result in increased genetic diversity with inbreeding: assortative overdominance generated by deleterious recessive alleles linked in repulsion phase or heterozygote advantage generated by linked or unlinked overdominant loci (Ohta and Kimura 1970, Palsson and Pamilo 1999). Linkage disequilibrium was low and similar under trioecy and androdioecy and therefore close physical linkage among deleterious recessives is unlikely to have generated much associative overdominance – but see (Szulkin et al. 2010, Chelo and Teotónio 2013, Noble et al. In Press) –. Identity disequilibrium was, however, important, particularly under androdioecy (Figure 3). If identity disequilibrium reveals associations of heterozygous genotypes across the genome, then the high genetic diversity that was maintained under androdioecy, during experimental evolution and after inbreeding (Figures 2 and 4), could well have been caused by overdominant loci.

Identity disequilibria caused by selfing should, however, reveal associations in homozygous genotypes (Weir and Cockerham 1973). Recent theoretical studies have shown that these identity disequilibria could maintain inbreeding depression by preventing purging of physically unlinked deleterious recessive alleles, and possibly by increasing Hill-Robertson effects and selective interference (Kamran-Disfani and Agrawal 2014, Roze 2015). Consistent with these ideas, androdioecious populations showed appreciable identity disequilibria (Figure 3), maintained genetic diversity during experimental evolution and after enforced inbreeding (Figures 2 and 4), and fertility, a fitness component (Carvalho et al. 2014b), did not evolve (Supplementary Figure 3).

If maintenance of genetic diversity during experimental evolution resulted from overdominant loci or deleterious recessives in identity linkage disequilibria, then androdioecious populations should have maintained ancestral inbreeding depression, but they did not (Figure 1). Since we assume that there was little mutation accumulation during the experiment, the simplest solution to this puzzle is to invoke that unlinked deleterious recessives were purged because of selfing, allowing then for the expression of a few unlinked overdominant loci that when in homozygosity resulted in high mortality (Nordborg et al. 1996). Note that recessive lethals in repulsion, even if unlinked, are effectively overdominant, cf. (Phillips and Johnson 1998). This would explain maintenance of genetic diversity under trioecy and androdioecy, but lack of inbreeding depression under androdioecy and monoecy.

A non-exclusive explanation would be the existence of overdominant epistasis for fitness, particularly negative dominance-by-dominance epistasis where homozygous combinations would similarly result in high mortality, as we have previously suggested to be the case during laboratory domestication under androdioecy (Chelo and Teotónio 2013). However, we recently described by genome-wide association analysis that the heritability of fertility is in large part determined by large effect additive-by-additive epistatic loci, and analysis of the genome sequence of the inbred lines used here suggests that the original wild genotypes hybridized to construct our populations are rarely if ever in repulsion phase (Noble et al. In Press). Both of these findings suggest a minor role for dominance-by-dominance epistasis is generating negative linkage disequilibrium. Until fitness loci can be mapped the jury will still be out regarding the role of dominance-by-dominance epistasis in the experimental evolution reproduction systems.

Monoecious populations reproduced exclusively by selfing and consequently hermaphrodites were homozygous throughout their genomes. It is then intriguing that in these populations genetic diversity after inbreeding was lower than that expected by chance (Figure 4D and E). One reason for this is the occurrence of selective interference between similarly fit genotypes during experimental evolution, though lack of a negative correlation between SNP differentiation during experimental evolution and inbreeding suggests otherwise (Figure 4C, Supplementary Figure 5). Alternatively, frequency-dependent selection where the survival of focal genotypes depends on other segregating genotypes could explain the maintenance of genetic diversity under exclusive selfing during experimental evolution but loss upon inbreeding. Similar phenomena have been observed in microbial evolution experiments (Ribeck and Lenski 2015), and also by us among two specific inbred lines derived from the lab adapted ancestor population (Chelo et al. 2013b).

In summary, partial selfing populations under androdioecy appear to gain the short-term benefits of selfing in purging deleterious genetic diversity and may reap the benefits of outcrossing in maintaining genetic diversity by overdominant loci that may be important for future adaptation. We find little evidence that linkage and identity disequilibria, or Hill-Robertson effects, have much bearing under partial selfing, at the population sizes employed. It remains to be shown whether partial selfing under other reproduction systems, particularly when hermaphrodites can outcross with each other, can also maintain genetic diversity without suffering from inbreeding depression (Winn et al. 2011, Noel et al. 2017). If such is the case, then the occurrence of mixed reproduction modes within natural populations is a step closer of being better understood.

## Acknowledgements

We thank F. Melo and A. Silva for help with the fertility assays; J. Costa, T. Guzella and L. Noble for support with the genotype analysis; and T. Guzella, L. Noble, M. Rockman and D. Roze for discussion. This work was partially supported by Fundação para a Ciência e Tecnologia through the FCT Investigator Programme (IF/00031/2013) to I.M.C.; the National Science Foundation (EF-1137835) to S.R.P., and the Human Frontiers Science Program (RGP0045/2010), the European Research Council (FP7/2007-2013/243285) and Agence Nationale de la Recherche (ANR-14-ACHN-0032-01) to H.T.

## Data Accessibility

Fertility and genotype data for the inbred lines associated with (Noble et al. In Press) has been archived in *FigShare* 10.6084/m9.figshare.5326777. All data and R code for analysis can be found in *FigShare* accessions: 10.6084/m9.figshare.5539606 (readme), 10.6084/m9.figshare.5539612 (code), 10.6084/m9.figshare.5539615 (dataset phenotypes and SNP genotypes), 10.6084/m9.figshare.5539618 (dataset haplotypes), and 10.6084/m9.figshare.5539627 (dataset simulated haplotypes).

## Supplementary Figures

**Supplementary Figure 1.**
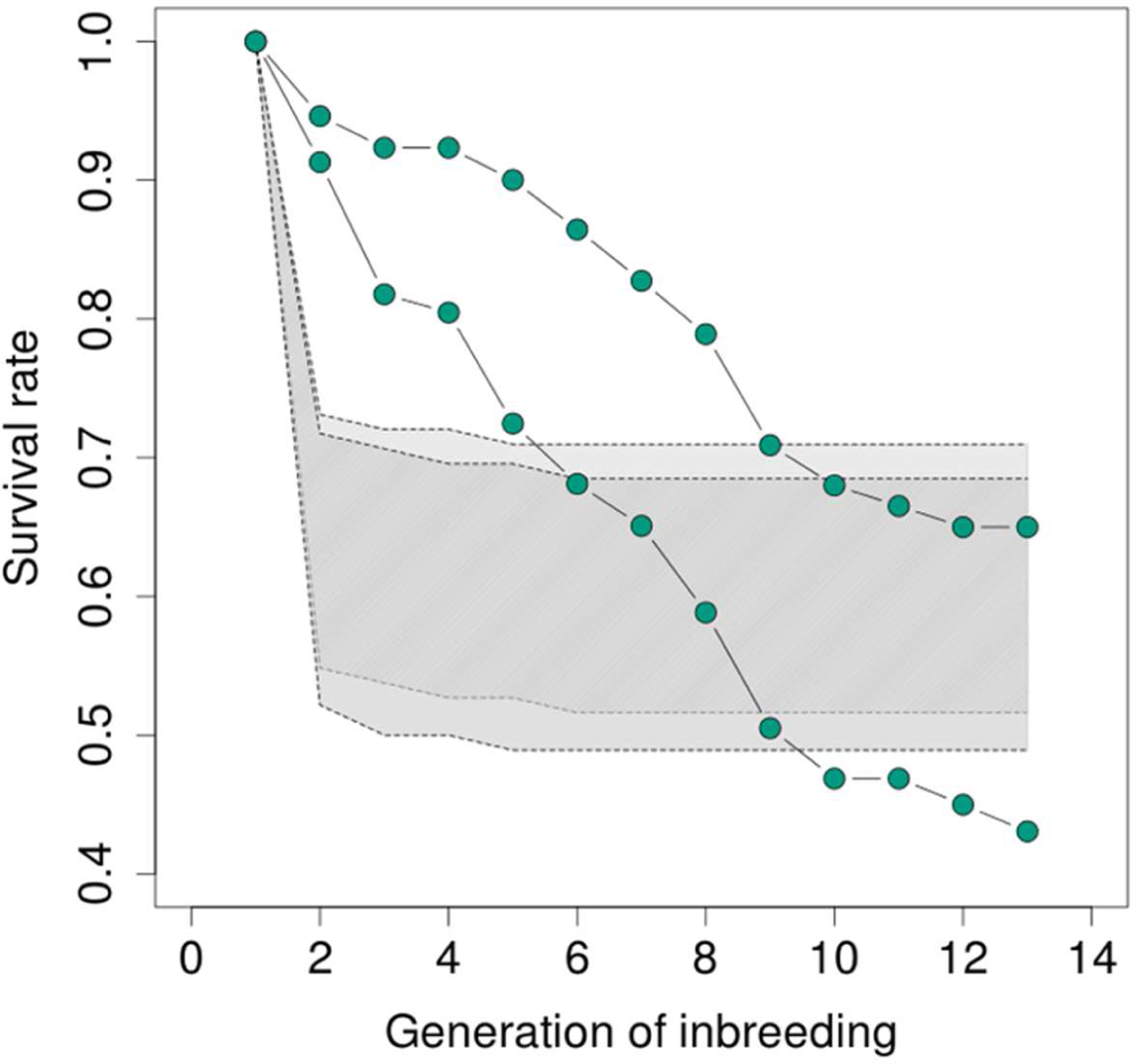
Expected proportion of lineage extinctions with generation of inbreeding by selfing of trioecious populations when females and/or heterozygous hermaphrodites for *fog-2(q71)* are sampled. Shaded region indicates the 95% confidence obtained with 1,000 simulations of the inbred line derivation. *fog-2* allele dynamics can be described as those of recessive lethal alleles, which are rapidly extinct. Comparison with observed survival rates (lines connecting green dots) reveal distinct dynamics from those obtained by this sampling process alone.

**Supplementary Figure 2.**
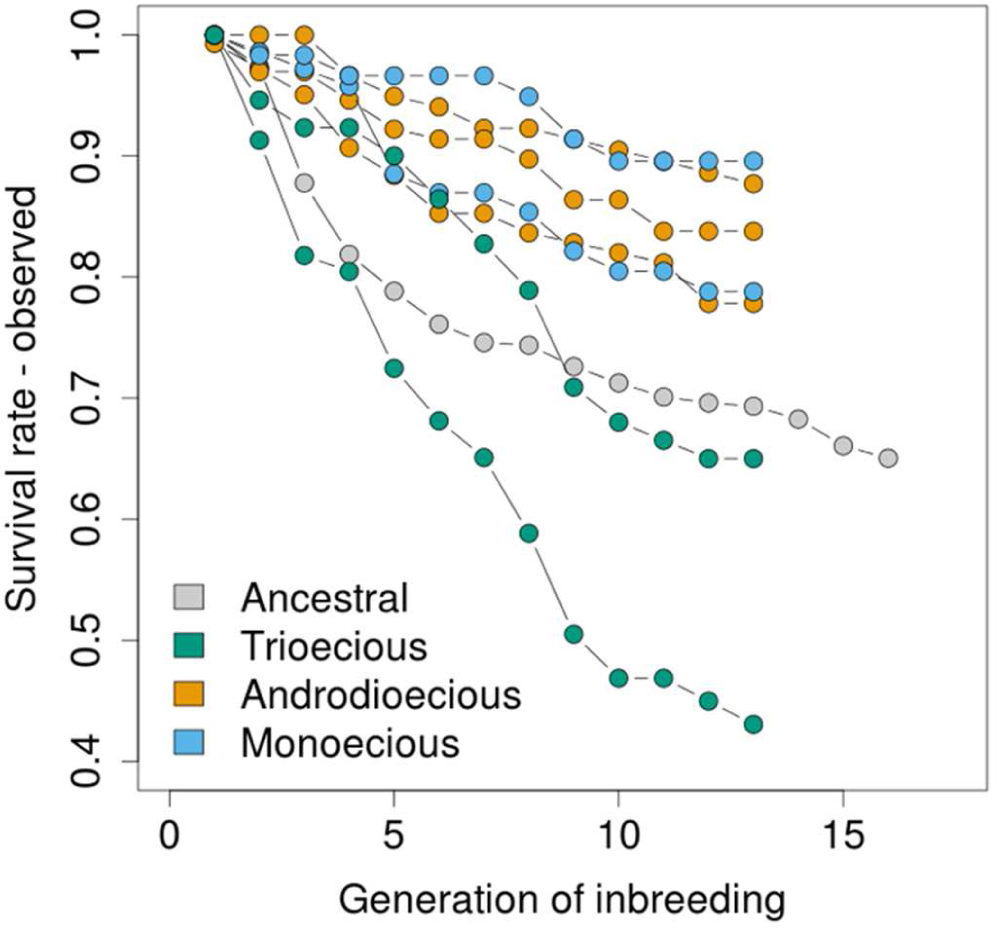
Observed proportion of lineage extinctions with generation of inbreeding by selfing. Each colored line represents a different replicate population from the three reproduction systems, and the laboratory adapted ancestral population.

**Supplementary Figure 3.**
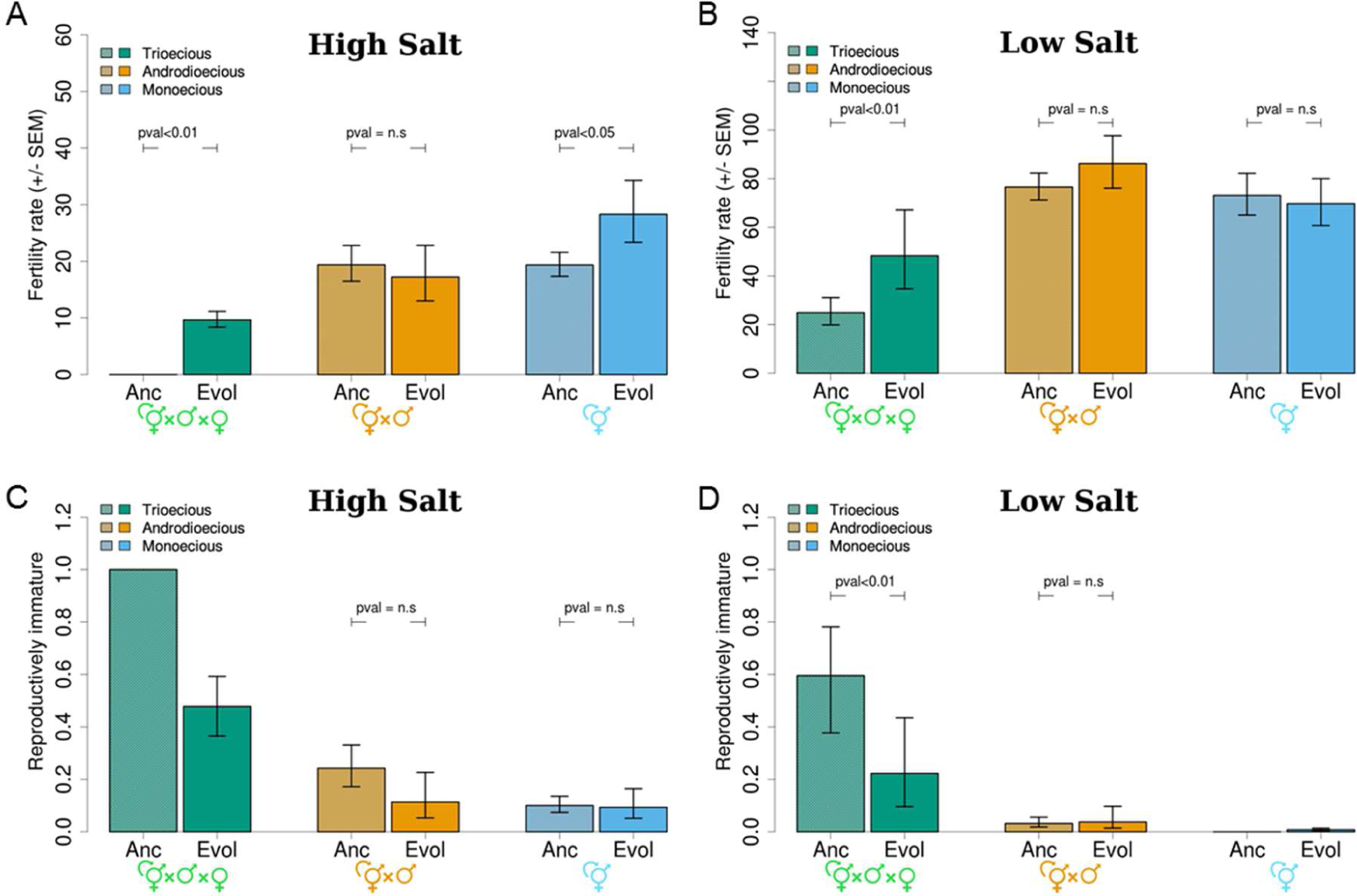
Fertility is modeled by dividing the number of first stage larvae (L1s) produced by each hermaphrodite/female into a class following a negative binomial distribution (panels A and B) and another class with no offspring (panels C and D). Individuals assigned to the “zero-class” are considered to be reproductively immature, or in the case of trioecy unmated females. Fertility values are given by the negative binomial distribution. Means and one standard error of the mean among replicates are shown in 305 mM NaCl (panels A and C) and 25 mM NaCl (panels B and D). Note that all individuals in the ancestral trioecious populations failed to produce any offspring in the high salt environment and, as such, individual assignment into a “zero-class” and a variable class is not adequate. See (Theologidis et al. 2014) for further details about adaptation under the three different reproduction systems.

**Supplementary Figure 4.**
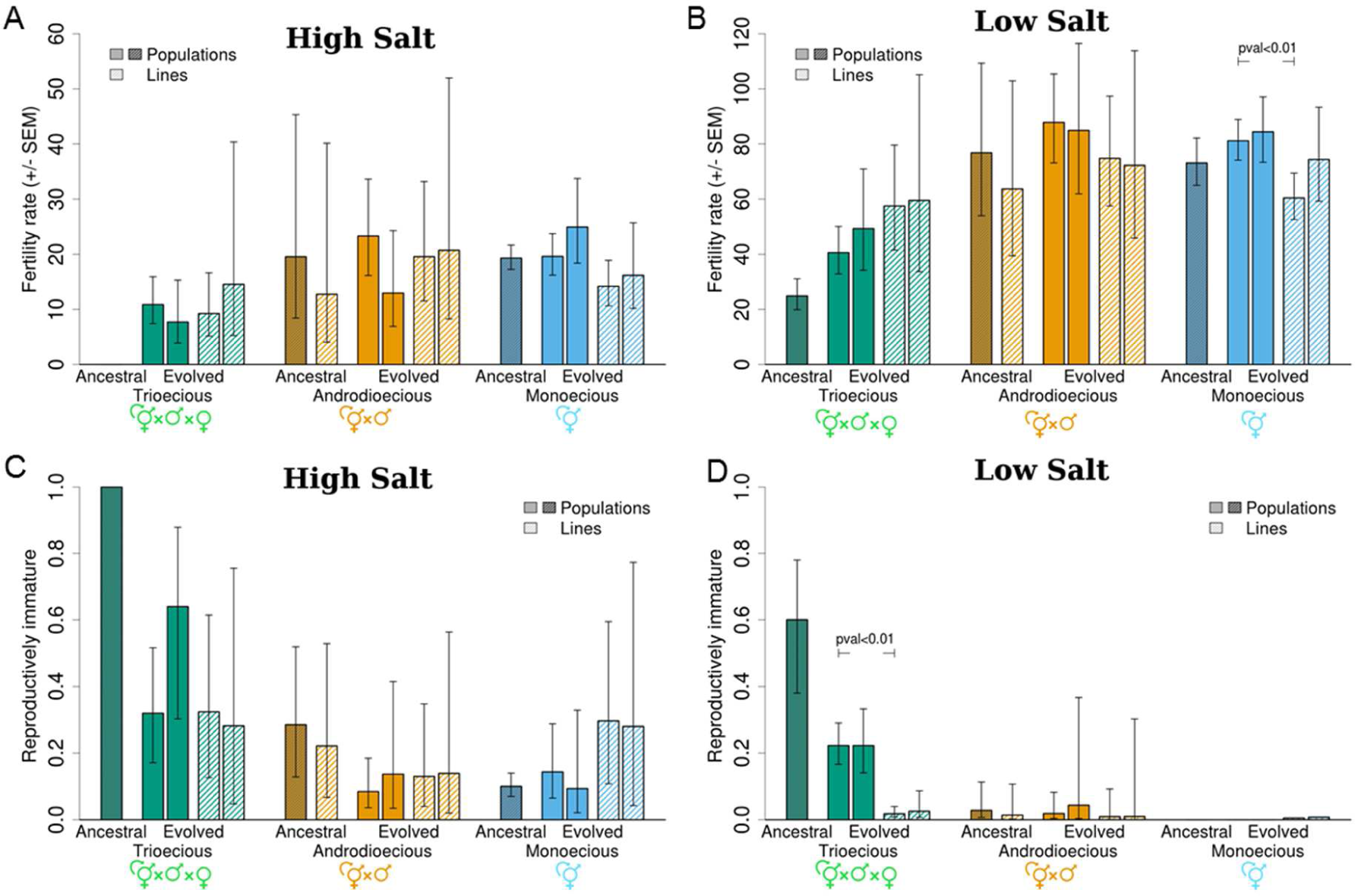
Mean fertility and one standard mean error among replicates of populations under different reproduction systems and among the inbred lines derived from them, in 305 mM NaCl (A) and 25 mM NaCl. Similarly, proportion of immature individuals or immature trioecious females at high (C) and low (D) salt conditions. The ancestral trioecious and ancestral monoecious populations, from which no inbred line derivation was done, were not included in this analysis and are shown only for comparative purposes.

**Supplementary Figure 5.**
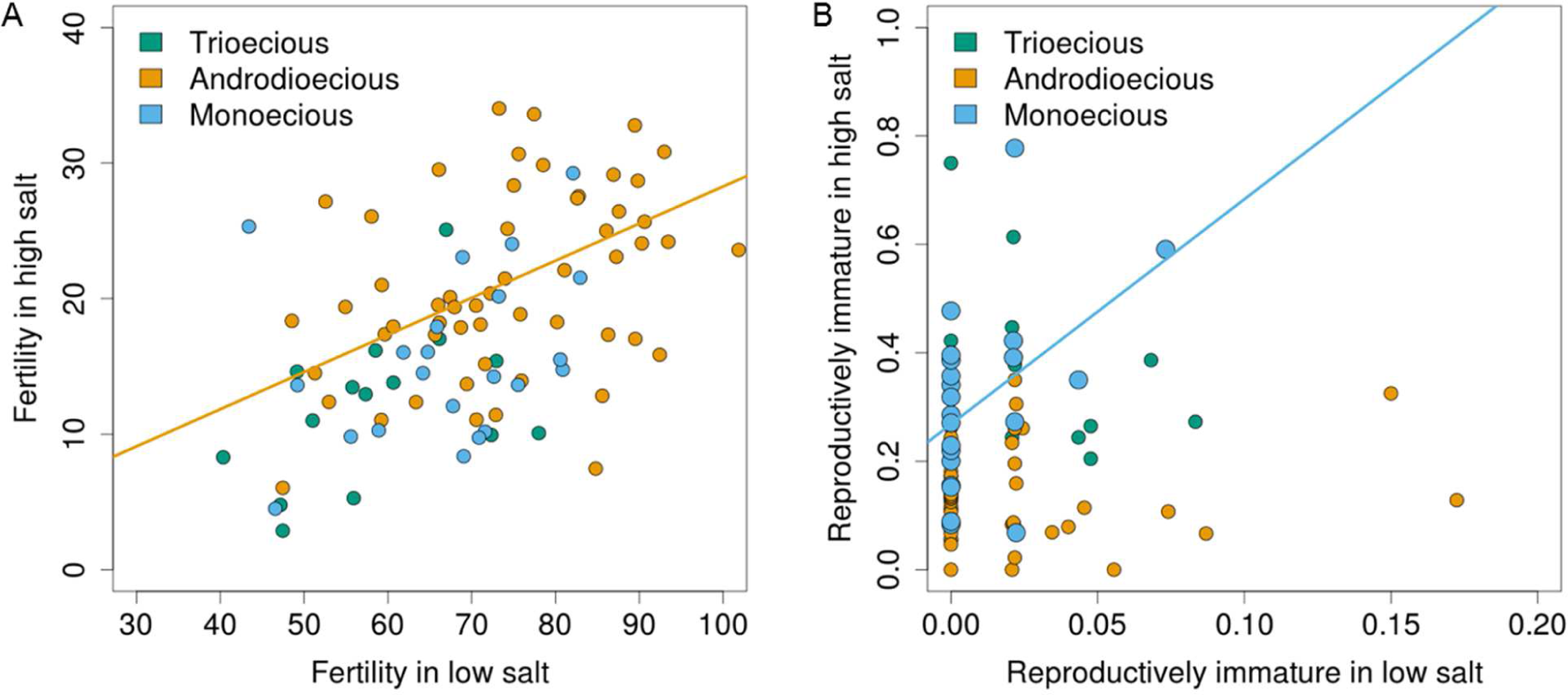
Fertility (A) or proportion of reproductively immature individuals (B) among the inbred lines are shown in high and low salt conditions. Significant genetic correlations between salt conditions are illustrated by the lines, as assessed with Spearman rank tests.

